# Lipid Droplet-Anchored Mitochondria Are More Sensitive to Cold in Brown Adipocytes

**DOI:** 10.1101/2020.04.07.029645

**Authors:** Mirza Ahmed Hammad, Liujuan Cui, Shuyan Zhang, Pingsheng Liu

**Affiliations:** National Laboratory of Biomacromolecules, CAS Center for Excellence in Biomacromolecules, Institute of Biophysics, Chinese Academy of Sciences, Beijing 100101, China; University of Chinese Academy of Sciences, Beijing 100049, China; School of Life Sciences, University of Science and Technology of China, Hefei, Anhui 230027, China

**Keywords:** Lipid droplet-anchored mitochondria, BAT, comparative proteomics, FA β-oxidation

## Abstract

Brown adipose tissue (BAT) are specialized for uncoupled heat production through mitochondria fueled majorly from fatty acids (FA) of lipid droplets (LDs). How the interaction between the two organelles contributes the generation of heat remains elusive. Here we report that LD-anchored mitochondria (LDAM) were observed in BAT of mice raised at three different temperatures, 30°C, 23°C, and 6°C. The biochemical analyses including Western blotting and electron transport chain subunits showed that LDAM were functional at given temperatures. Comparative proteomics analysis was conducted and revealed that these LDAM had protein level differences from cytoplasmic mitochondria (CM) at different temperatures. Higher expressions of proteins at low temperature were observed for i) FA β-oxidation in LDAM including FA synthesis, and uncoupling, ii) pseudo-futile cycle in CM, and iii) two shuttle systems; glycerol 3-phosphate in both CM and LDAM, and citrate malate in CM. Together, these results suggest that LDs and LDAM are a preorganized and functional organelle complex that permits the rapid response to cold environment.

## Introduction

Brown adipose tissue (BAT) is enriched in lipid droplets (LDs) and mitochondria, and specialized in non-shivering thermogenesis through adaptive uncoupling process. Fatty acids (FAs) majorly released from LDs are primary source of energy for non-shivering thermogenesis, with glucose acting as an additional source when required.

LD-mitochondrial association has been observed in muscle cells where degree of contact was enhanced in response to exercise (Tarnopolsky et al., 2007). In cardiomyocytes Perilipin 5 (PLIN5) was found to play an important role in this interaction (Wang et al., 2011). It was also observed that this contact was dependent on microtubule detyrosination (Herms et al., 2015) and enhanced by Mitofusin 2 in BAT (Boutant et al., 2017). The processes of triacylglycerol (TAG) hydrolysis and FA β-oxidation are crucial to the maintenance of non-shivering thermogenesis in BAT whereby direct physical contact between the organelles can greatly facilitate this transport.

Latest studies evidenced physical contact between LDs and mitochondria in BAT, and identified PLIN1 and PLIN5 as major factors for the contact (Olzmann and Carvalho, 2018). At least two contact forms have been reported. One study showed that BAT LDs were tight associated with mitochondria (Yu et al., 2015) and later study identified that these two organelles cannot be separated by ultracentrifugation and named this population of mitochondria as LD-anchored mitochondria (LDAM) (Cui et al., 2019). In addition, this non-strippable contact is also identified in skeletal muscle and heart, but not liver in mouse and monkey (Cui et al., 2019). Another study reported that half of LD-associated mitochondria can be stripped by 9,000*g* centrifugation and termed the mitochondrial fraction as peridroplet mitochondria (PDM) (Benador et al., 2018). PDM are exhibited distinct pathway activation from cytoplasmic mitochondria (CM), while the pathway activation for LDAM remains elusive. The fact that PDM are strippable and LDAM are not suggests that with respect to LD association, different mitochondrial subpopulations act differently based on strength of association.

Therefore, it is important to understand pathway level changes between different mitochondrial subpopulations. In a recent proteomics study, Sanchez-Gurmaches et. al. (Sanchez-Gurmaches et al., 2017) observed a paradoxical behavior of FA β-oxidation and FA recycling in BAT proteome at low temperature. The first evidence of FA recycling dates back to 1960 when conversion of extra pyruvate into glucose was observed to be overtaken by high production of labelled glycerol in a process termed as glyceroneogenesis (Ballard et al., 1967). In this study, we isolated LD and CM fractions from mice housed at thermoneutral, room, and low temperatures, and then pursued differences between LDAM and CM using a comparative proteomic method. We observed that LDAM were a different mitochondrial subpopulation with unique protein composition, especially the upregulated proteins for heat production in LDAM under low temperature, suggesting that LDAM were more sensitive than CM in response to cold. Additionally, deep pathway analysis indicated enhanced expression of proteins leading to a possibility of a pseudo-futile cycle with two shuttle systems.

## Experimental Procedure

### Materials

The TRIzol reagent, HCS LipidTOX™ Green Neutral Lipid Stain (H34475), HCS LipidTOX™ Red Neutral Lipid Stain (H34476), MitoTracker™ Red CMXRos (M7512), PageRuler prestained protein ladder, and SYBR SELECT MASTER MIX were from ThermoFisher SCIENTIFIC. Potassium ferrocyanide was from Sigma-Aldrich. 25% Glutaraldehyde solution, EMbed 812 kit, Uranyl acetate and Lead citrate were purchased from Electron Microscopy Sciences. Osmium tetraoxide (EM grade) was from Nakalai Tesque. Anti-Perilipin1 was a gift from Dr. Guoheng Xu. Anti-ATGL was from Cell Signaling Technology. Antibodies against Tim23 and OPA1 were from BD Biosciences. Antibodies against Rab18, and ACSL5 were from Abclonal Technology. Antibodies against p-HSL and Mfn2 were from Bioworld Technology. Antibodies against ADRP, total OXPHOS, and UCP1 were from Abcam. Anti-SOD2 antibody was obtained from Proteintech. Anti-VDAC and anti-Prohibitin antibodies were obtained from Millipore. Western Lighting Plus ECL was from PerkinElmer.

### Animal Studies

Eight-week-old male C57BL/6 mice were purchased from Vital River Laboratories (VLR) (Beijing). Chow diet was purchased from Beijing Keaoxieli Feed Co., Ltd. The animals were maintained on 12 h:12 h light-dark cycles. The mice were randomly divided into 3 groups, and were then raised in incubators at different temperatures (30°C, 23°C, 6°C). The mice were allowed free access to standard rodent chow and water ad libitum. All animal protocols were approved by the Animal Care and Use Committee of the Institute of Biophysics and University of Chinese Academy of Sciences under the permission number SYXK (SYXK2016-11).

### Isolation of LDs from Mouse Brown Adipose Tissue and LipidTOX Staining

LDs were isolated by a modified method (Ding et al., 2013; Liu et al., 2004; Yu et al., 2015). Since brown adipocytes have very large LDs with little cytosol, we used a low speed centrifugation to isolate LDs from brown adipose tissue (BAT). First, we took the interscapular BAT from C57BL/6 male mice into ice-cold saline with 0.5 mM PMSF and discarded any attached white fat tissue (WAT). Then the BAT was cut into 1-2 mm^3^ pieces and transferred into 2.5 mL ice-cold Buffer A (20 mM Tricine, pH 7.6, 250 mM Sucrose) with 0.5 mM PMSF. The minced BAT was homogenized by passing through a stainless steel sieve (200 mesh, 70 micron), the whole cell lysate (WCL) was collected and incubated on ice for 20 min. Then the WCL was centrifuged at 2,000*g* (Low speed, L) for 6 min at 4°C. The LD fraction on the top was collected into a new 1.5 mL microcentrifuge tube. The remaining supernatant was subjected to ultracentrifugation (Optima™ Ultracentrifuge TLA 100.3) at 303,475*g* (High speed, H) for 15 min. After the centrifugation, the middle clear supernatant was collected as cytosol (Cyto) and the pellet was collected as total membrane (TM). LDs were washed with 200 μL Buffer B (20 mM HEPES, pH 7.4, 100 mM KCl, 2 mM MgCl_2_) three times. Finally, 800 μL acetone and 300 μL chloroform was added to the LDs followed by thoroughly vortexing. Then the mixture was centrifuged at 21,130*g* for 10 min, the pellet was LD proteins. The LD proteins were subjected to proteomics analysis or were dissolved with 2 × SDS-sample buffer and denatured at 95°C for 5 min for silver staining and Western blotting analysis.

Isolated LDs were resuspended with Buffer B and stained with LipidTOX Green/Red on ice for 20-30 min. The ratio of LipidTOX to LD sample was 1:500 (v/v). The LDs were then visualized using an Olympus FV1200 Imaging System. Intact LDs were determined by their spherical shape and smooth edges in micrographs from both fluorescence microscopy and electron microscopy (EM).

### Mitochondrial Isolation from Mouse Brown Adipose Tissue and MitoTracker Staining

Mitochondria were isolated by a modified method (Pu et al., 2011; Yu et al., 2015). WCL was centrifuged at 500*g* for 10 min to remove the nucleus and the supernatant was collected as the post-nuclear supernatant (PNS). The PNS was centrifuged at 8,000*g* for 10 min at 4°C. The pellet was resuspended and washed twice with 500 μL Buffer B as rough mitochondria. The mitochondrial fraction was applied carefully to the top of a Percoll step gradient (3 mL 50% and 8 mL 25%) and was centrifuged at 41,000*g* (Optima™ L-100 XP Ultracentrifuge) for 45 min at 4°C. The interface between 50% and 25% Percoll was collected 200 μL into each tube. The collected fraction was diluted by the addition of 1 mL Buffer B, vortexed thoroughly, and centrifuged at 21,130*g* for 10 min. The pellet was washed three times with 1mL Buffer B to remove Percoll.

Isolated mitochondria were resuspended with Buffer B and stained with MitoTracker Red on ice for 20 min, with a ratio of MitoTracker:Mitochondria of 1:1,000 (v/v). The samples were visualized with an Olympus FV1200 Imaging System. Intact mitochondria were determined by their spherical shape and smoothly edges in micrographs from both fluorescence microscopy and EM.

### Electron Microscopy

Through ultra-thin sectioning, the ultra-structure of BAT was examined by transmission electron microscopy (TEM). Briefly, after collected and rinsed, the BAT tissue was cut into small pieces (about 1 mm^3^). These pieces were fixed in 2.5% (w/v) glutaraldehyde in 0.1 M PB (pH 7.2) for 2 h at room temperature. Subsequently, they were fixed in 1% (w/v) osmium tetraoxide (with 1% potassium ferrocyanide) for 2 h. After dehydrated in an ethanol series at room temperature, the samples were embedded in EMbed 812 and finally prepared as 70 nm ultra-thin sections. After stained with uranyl acetate and subsequently with lead citrate at room temperature, the sections were observed with Tecnai Spirit electron microscope.

The isolated LDs and mitochondria were also examined by TEM. Briefly, isolated LDs were embedded in 4% agarose first. After solidification, the samples were cut into small blocks (∼1 mm^3^). The pellets of isolated mitochondria were fixed directly. The samples were prefixed in 1% glutaraldehyde for 30 min and then post-fixed in 1% osmium tetraoxide (with 1% potassium ferrocyanide) for 30 min at room temperature. The samples were finally prepared as 70 nm sections and viewed with electron microscope.

For cryo-EM analysis, 4 μL of isolated mitochondria was placed onto grids and blotted for 3 s in 100% humidity using a FEI Vitrobot Mark IV. Then the grids were vitrified by quickly plunging into pre-cooled ethane pot within the liquid nitrogen. Micrographs were recorded with an FEI 300-kV Titan Krios cryo-electron microscope equipped with a Gatan UltraScan4000 (model 895) 16-megapixel CCD.

### Silver Staining and Western Blotting

For the silver staining, the gel was fixed in fixation buffer for 30 min and was incubated in sensitizer buffer at room temperature for 30 min. The gel then was washed four times with ddH_2_O for 5 min each time. The washed gel was treated with silver staining buffer at room temperature for 20 min, followed by developing buffer until the bands appeared. The reaction was blocked in stopping buffer immediately after achieving satisfactory resolution.

For Western blotting, proteins from different fractions were separated by sodium dodecyl sulfate-polyacrylamide gel electrophoresis (SDS-PAGE) and were transferred to PVDF membranes. The membranes were blocked with 5% BSA at room temperature for 1 h and were then incubated with primary antibodies at room temperature for 1 h or 4°C overnight. The membranes were washed 3 times with washing buffer for 5 min each time followed by incubation in the indicated secondary antibodies at room temperature for 1 h. The protein bands were visualized with Western Lighting Plus ECL.

### Quantitative Real-Time PCR

Total RNA was extracted with TRIzol reagent and reverse transcribed into cDNA according to the manufacturer’s protocol (Promega and TAKARA). qRT-PCR was performed using SYBR SELECT MASTER MIX and Biosystems QuantStudio 7 Flex for 40 cycles and the fold change for all the samples was calculated by the 2^-ΔΔCt^ method, n ≥ 6. β-actin was used as housekeeping gene for mRNA expression analysis.

### Sample Preparation for Proteomic Study

The extracted proteins from LDs or mitochondria were reduced with 10 mM dithiothreitol for 2 h at room temperature and then immediately alkylated with 20 mM iodoacetamide for 45 min at room temperature in the dark. The resulting protein solution was diluted with 100 mM ammonium bicarbonate to less than 2 M urea and digested with trypsin (50:1) at 37°C overnight. After quenched with formic acid, the peptide mixtures were desalted followed by vacuum centrifugation. The dried peptides were dissolved with 100 mM triethylammonium bicarbonate and then labeled with 6-plex Tandem Mass Tags (TMT) reagents (Thermo Fisher Scientific) according to the manufacturer’s instructions. The samples were labeled as follows: TMT-126/-127 for samples at 30°C, TMT-128/-129 for samples at 23°C and TMT-130/-131 for samples at 6°C. For mitochondrial samples, prior to MS analysis, the mixed TMT-labeled samples were further fractionated using a Waters XBridge BEH130 C18 column (4.6 × 250 mm, 5 μm particles) in an L-3000 HPLC System (Rigol). All fractions were collected at 90 s intervals and concatenated into 10 fractions and lyophilized.

### LC-MS/MS Analysis

Nano LC-MS/MS experiments were performed on a Q Exactive (Thermo Fisher Scientific) equipped with an Easy n-LC 1000 HPLC system (Thermo Fisher Scientific). The above labeled peptides were loaded onto a 100-μm id × 2-cm fused silica trap column packed in-house with reversed phase silica (Reprosil-Pur C18-AQ, 5 μm, Dr. Maisch GmbH, Germany) and then separated on a 75-μm id × 20-cm C18 column packed with reversed phase silica (Reprosil-Pur C18-AQ, 3 μm, Dr. Maisch GmbH, Germany). The loaded peptides on the column were eluted with a 78-min linear gradient. The solvent A consisted of 0.1% formic acid in water solution and the solvent B consisted of 0.1% formic acid in acetonitrile solution. The following segmented gradient was set at a flow rate of 300 nl/min: 5–8% B, 8 min; 8–22% B, 50 min; 22– 32% B, 12 min; 32-95% B, 1 min; 95% B, 7min.

The MS analysis was performed with Q Exactive mass spectrometer (Thermo Fisher Scientific). In a data-dependent acquisition mode, the MS data were acquired in the Orbitrap at a high resolution 70,000 (m/z 200) across a mass range of 300–1600 m/z. The target value was 3.00E+06 with a maximum injection time of 60 ms. The top 20 most intense precursor ions were selected from each MS full scan with isolation width of 2 m/z for fragmentation in the HCD collision cell with normalized collision energy of 30%. Subsequently, MS/MS spectra were acquired in the Orbitrap at a resolution of 17,500 at m/z 200. The target value was 5.00E+04 with a maximum injection time of 80 ms. The dynamic exclusion time was 40 s. The nano electrospray ion source settings were as follows: the spray voltage was 2.0 kV, no sheath gas flow, and a heated capillary temperature of 320°C.

### Protein Identification and Quantification

The raw MS data from Q Exactive were analyzed with Proteome Discovery (version 2.2.0.388, Thermo Fisher Scientific) using Sequest HT search engine for protein identification and Percolator for FDR (false discovery rate) analysis. The UniProt mouse protein database (updated on 10-2017) was used for searching the data. Searching parameters were set as follows: trypsin was selected as enzyme and two missed cleavages were allowed for searching; the mass tolerance of precursor was set as 10 ppm and the product ions tolerance was 0.02 Da; TMT 6plex (lysine and N-terminus of peptides) and cysteine carbamidomethylation were specified as fixed modifications; the methionine oxidation was chosen as variable modification. FDR analysis was performed using Percolator and FDR <1% was set for protein identification. The peptide confidence was set as high for peptide filter. Protein quantification was also performed on Proteome Discovery 2.2.0.388 using the ratio of the intensity of reporter ions from the MS/MS spectra. Only unique and razor peptides of proteins were selected for protein relative quantification. The co-isolation threshold was specified as 50% and average reporter S/N value should be above 10. The normalization to the protein median of each sample was used to correct experimental bias and the normalization mode was selected as total peptide amount.

### Experimental Design and MS Data Analysis

In this study, we used isolated LDs and mitochondria from BAT of mice raised at different temperatures for quantitative proteome to find out differences in their response to temperature. Thus, two groups of mice (R1 and R2, i.e., two biological replicates), 4 mice each, were raised for one month at 30°C, 23°C, and 6°C respectively. BATs from one group (four mice) at each temperature were pooled as one sample, and subjected to isolation of LD and mitochondria. The 6-plex TMT reagents were used for quantitative proteomic study on LD fraction or mitochondrial fraction at three temperatures respectively.

A commutative approach was used to analyze the proteomes. In step 1, proteins with a peptide number < 5 were removed from analysis. In step 2, the proteins which did not show a reproducible trend of expression change across all three temperatures in both R1 and R2 groups were removed from analysis. Step 3 was based on expression rates. Any protein fulfilling criteria of (highest expression rate (from three temperatures) – lowest expression rate (from three temperatures) < 20 for CM) and (highest expression (from three temperatures) – lowest expression (from three temperatures) < 15 for LD) were considered as proteins showing no change in expression. For the comparison of protein expressions between any two temperatures, proteins meeting the criteria [highest expression rate (from two temperatures) – lowest expression rate (from two temperatures) < 15 for CM)] and (highest expression (from two temperatures) – lowest expression (from two temperatures) < 10 for LD) were considered as showing no change. In step 4, the relative strength of expression (RSE) was calculated for proteins at given temperatures as

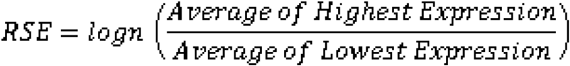

The RSE provided a better representation of change of expression at given temperature. The mitochondrial proteins were mapped using DAVID web-service and Mitocarta 2.0. For pathway analysis, primarily temperature based clustering was performed through the Perseus tool (Tyanova et al., 2016). Wiki pathway and KEGG databases were used for keywords-based pathway clustering. Heatmaps were drawn using Plotly.

## Results

### LDAM Exist in BAT of Mice Housed at 30°C, 23°C, and 6°C

Previously, LDAM was found and similar tight contact between LDs and mitochondria was also observed in skeletal muscle, heart, and BAT in both mouse and monkey (Cui et al., 2019). But why and how these mitochondria anchor on LDs remain unknown. Furthermore, most of previous studies about the two organelle contact were conducted either at 23°C or 4°C, while the thermoneutral temperature for mice is 29-30°C (Lodhi and Semenkovich, 2009). To investigate potential function of LDAM that respond to different temperatures, we raised mice at 30°C, 23°C, and 6°C, and then conducted indicated experiments on them (Fig. 1A). Preliminary studies on expression of genes suggested that reduced temperature was associated with a dramatic increase in RNA expression of *adrp* and *ucp1* (Fig. 1B). Western blotting (WB) further confirmed ADRP/PLIN2 and UCP1 as significantly cold-induced proteins (Fig. 1Cb). Increased expression of ADRP/PLIN2 with cold exposure was also observed in a previous study of mice housed at 4°C (Yu et al., 2015). Moreover, BAT activity-related genes *cgi58, cidec, cpt1b, pgc1α, rab5b*, and *rab18* also showed significantly enhanced expression as determined by RT-qPCR (Fig. 1B). H&E staining demonstrated that most brown adipocytes from the 30°C group were similar to white adipocytes with a unilocular phenotype (Haugen and Drevon, 2007) (Fig. 1Dd). In contrast, the cells of the 23°C and 6°C groups had multilocular LDs with reduced size (Barre et al., 1986) (Fig. 1De and f). Immunohistochemistry (Fig. 1Dg-i) results showed that dramatic increase in UCP1 expression was directly linked with low temperature, which agreed with WB data (Fig. 1Cb). Detailed intracellular structures of brown adipocytes examined by transmission electron microscopy (TEM) showed clear change in mitochondrial morphology from elongated at 30°C (Fig. 1Dj) to sphere-shape at 23°C and 6°C (Fig. 1Dk and l). While moving from 30°C to 23°C and 6°C, increased mitochondrial cristae were also observed. More important, the physical contact between LDs and mitochondria in BAT was also detected at 30°C (Fig. 1Dj, arrows) with no obvious differences in the amount of contact seen at the 23°C and 6°C (Fig. 1Dk and l, arrows).

**Figure 1.**
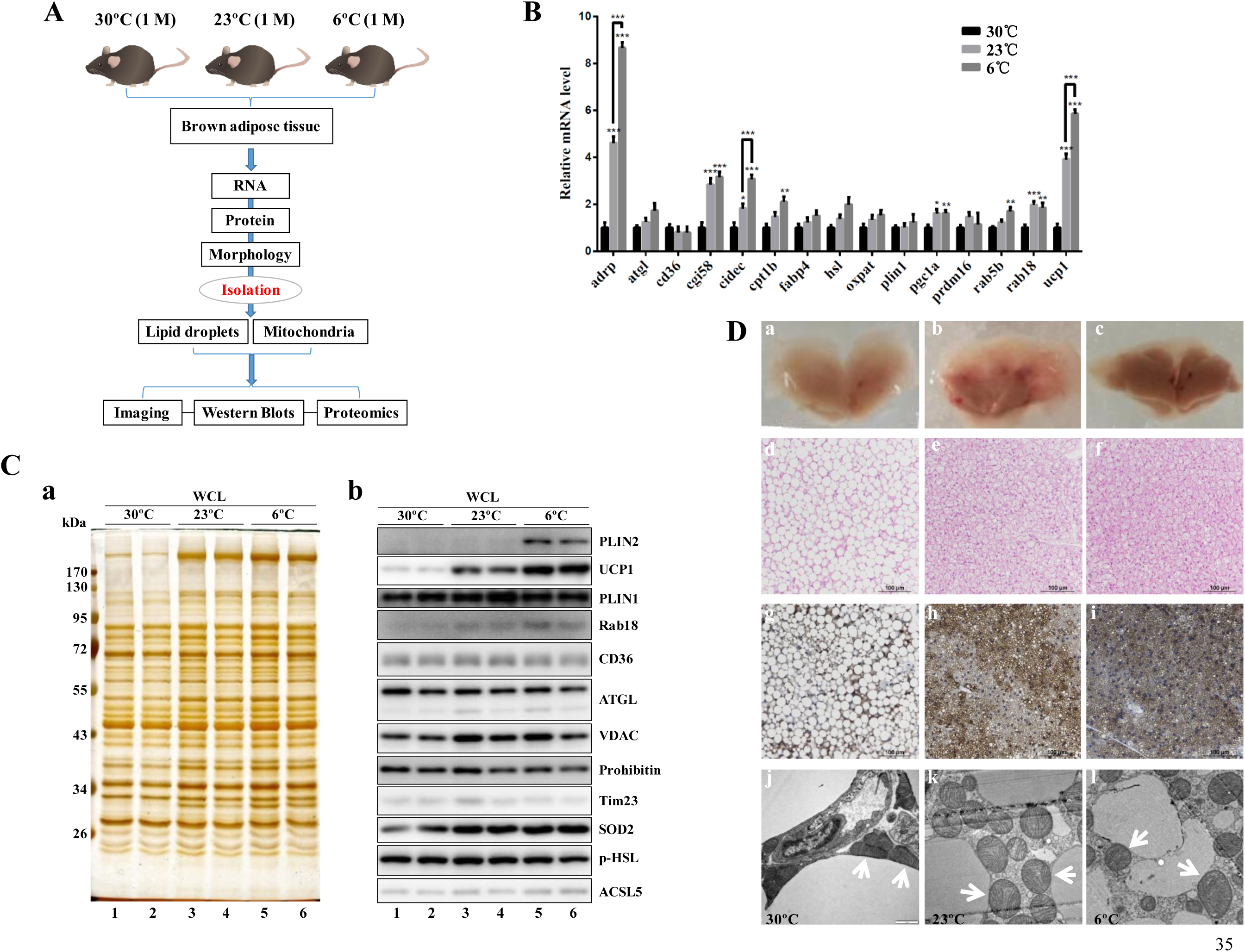
Contact between Lipid Droplets and Mitochondria in Mouse BAT from Three Temperatures. Eight-week-old male C57BL/6 mice (eight mice per group) raised at 23°C were transferred to incubators at different temperatures (30°C, 23°C, and 6°C) for one month. Then their interscapular brown adipose tissue (BAT) was collected and analyzed (A-D). **A** Schematic representation of the analysis of isolated lipid droplets (LDs) and mitochondria (CM) from mouse BAT. **B** Expression of metabolic-related genes in BAT using real-time PCR. n = 8. Statistical analysis was performed using Student’s t test. *, *P*<0.05; **, *P*<0.01; ***, *P*<0.001. **C** Protein profiles of mouse BAT using silver staining (left panel) and Western blotting of metabolic-related proteins (right panel). Each sample was from four mice. **D** Representative H&E staining (d, e, f), UCP1-immunohistochemistry (g, h, i), and transmission electron microscopy (TEM) (j, k, l) of BAT (a, b, c). Arrows point to the physical contact between LDs and mitochondria in BAT.

To further confirm this contact and to determine the effect of temperature on the contact, LDs and cytosolic mitochondria (CM) were isolated. First, LDs and CM were isolated from mice housed at 23°C and stained with LipidTOX and MitoTracker, respectively, to verify the quality of those isolated organelles. Both isolations were visualized through DIC (Fig. 2Aa and e) and fluorescence signal (Fig. 2Ab and f). The overlap of globe-shaped structures and LipidTOX positive structures in LD isolation confirmed that they were LDs (Fig. 2Aa and b). The overlap of MitoTracker signal and DIC structures in CM isolation proved isolated CM (Fig. 2Ae and f). Furthermore, CM isolation showed only mitochondrial structures as observed through TEM (Fig. 2Ag) and cryo-EM (Fig. 2Ah). TEM demonstrated mitochondria in physical contact with surface of the isolated LDs (Fig. 2Ac and d, arrows). The contact was further verified using biochemical means. The isolated LDs and CM shared similar protein compositions revealed by silver staining (Fig. 2Ba, lanes 1 and 2), while WB showed that LD fraction contained CM proteins including UCP1, Tim23, VDAC, and Mfn2 (Fig. 2Bb).

**Figure 2.**
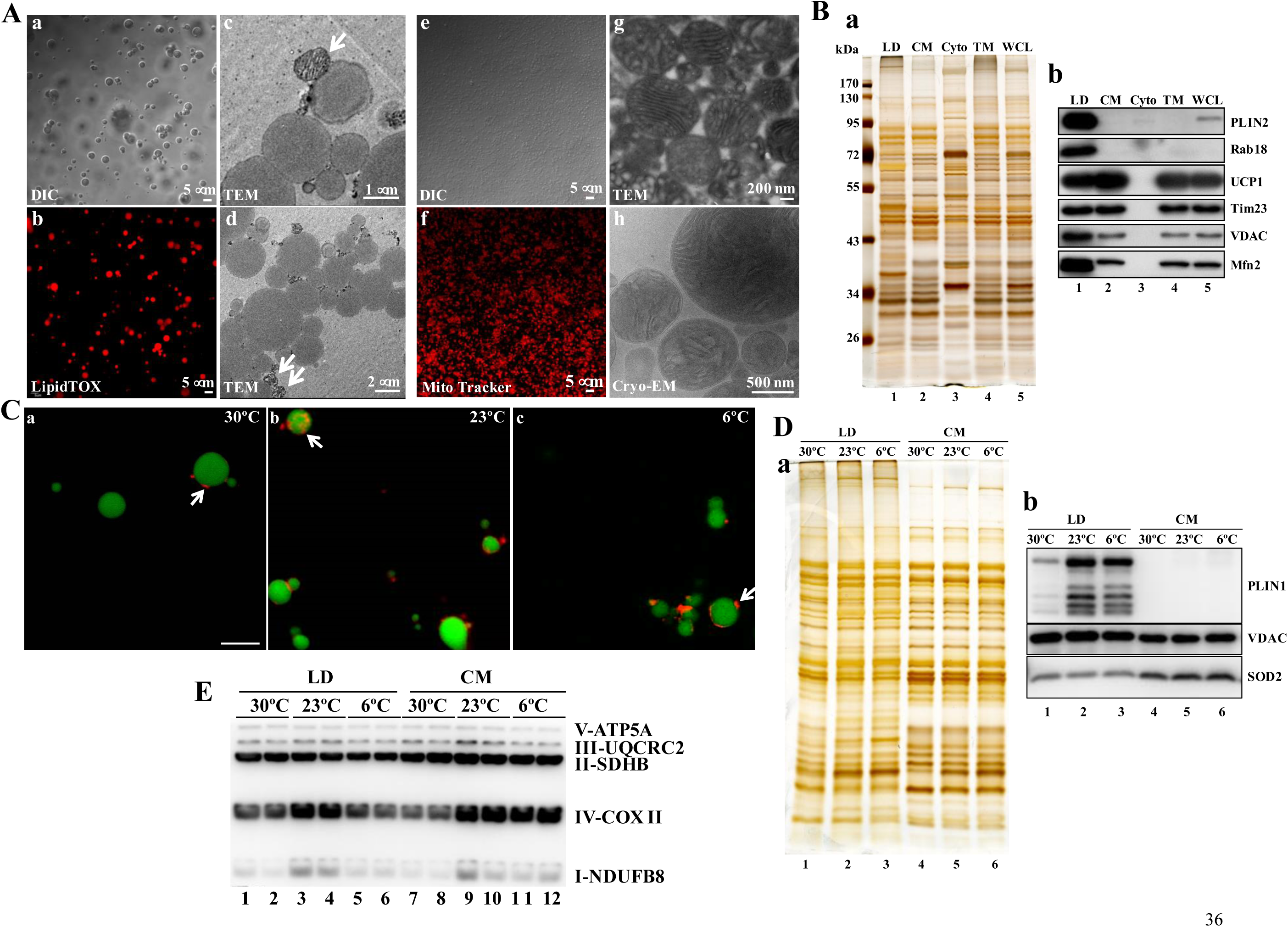
Anchoring Mitochondria to Lipid Droplets in Mouse BAT from Three Temperatures. **A** Morphologic analysis of isolated BAT LDs (a-d) and mitochondria (e-h) from mice under 23°C, including DIC (a, e), LipidTOX Red staining (b), MitoTracker Red staining (f), TEM (c, d and g), cryo-electron microscopy (Cryo-EM) (h). Arrows indicate LD-anchored mitochondrion in isolated LDs. **B** Isolated LDs, CM, TM, Cyto, and WCL were analyzed by silver staining (left panel), and Western blotting (right panel). TM, total membrane; Cyto, cytosol; WCL, whole cell lysate. **C** Isolated BAT LDs were double stained with LipidTOX (Green) for LDs and MitoTracker (Red) for mitochondria from different temperatures (30°C, 23°C, and 6°C). Arrows point to the physical contact between LDs and mitochondria in isolated BAT LDs after a merge of the signals. Bar = 5 μm. **D** Isolated BAT LDs and CM from different temperatures (30°C, 23°C, and 6°C) were analyzed by silver staining (left panel), and Western blotting (right panel). **E** Isolated LDs and CM were analyzed by Western blotting for the indicated mitochondrial OXPHOS complexes (I, II, III, IV and V) subunits.

Using this verified method, LDs and CM were isolated from mouse BAT housed at three temperatures. To determine the contact between them, the isolated LDs were stained with LipidTOX (green) for LDs and MitoTracker (red) for mitochondria (Fig. 2C). The MitoTracker red signals were detected on the LipidTOX green-stained sphere structures from all three temperatures (Fig. 2Ca-c, arrows). Then these isolations were further examined to confirm the contact of two organelles using silver staining and WB. Silver staining result showed that protein profiles had only minor differences among the sample from three temperatures for both LDs and CM (Fig. 2Da, lanes 1-3 and 4-6). In agreement with the previous study (Cui et al., 2019), the protein profile of LDs was similar to CM, which was further verified by WB results of mitochondrial proteins VDAC and SOD2 (Fig. 2Db).

Later on, electron transport chain (ETC) was analyzed in LDAM from LD fraction and CM from mitochondrial fraction at 30°C, 23°C, and 6°C. The amounts of proteins NDUFB8 from ETC subunit complex-I (C-I) and COX II from ETC subunit C-IV were elevated at 23°C in both LDAM and CM while SDHB, UQCRC2, and ATP5A from ETC subunits C-II, C-III, and C-V showed no observable change with varying temperature (Fig. 2E). These results again confirmed that the contact between LDs and mitochondria exists under three temperatures even at 30°C.

### Comparative Proteomics Analysis of Isolated Lipid Droplets and Mitochondria

A simple overlook of previously published proteomes indicates presence of mitochondrial proteins in LD proteomes (Table S1). It, therefore, also suggests that the LDs and mitochondria have tight contact in different oxidative tissues and in many eukaryotes. Still, evolutionary perspective of this contact may still need to be uncovered. In current study, with established temperature mouse model and verified isolation methods, the contact between LDs and mitochondria was able to be obtained for further biochemical study. Thus, two groups of mice (R1 and R2) were raised for one month at each of three temperatures, 30°C, 23°C, and 6°C. The two groups in each temperature consisted of 4 mice each. Their BATs were collected and BATs from four mice for each condition were pooled as one sample, and then subjected to LD and CM isolation. Before sending for proteomic analysis, their proteins were analyzed using silver staining and WB for quality control (Fig. 3A). First, silver staining result presented that protein compositions of two groups for each temperature were nearly identical and there were some differences among three temperatures (Fig. 3Aa). Second, WB result demonstrated that LD marker proteins PLIN1, PLIN2, and PLIN5 were only detected in LD fractions and increased significantly during temperature drop (Fig. 3Ab, lanes 1-6). Third, both silver staining and WB confirmed that LD fractions contained mitochondrial proteins, presenting that LDAM and CM were successfully obtained and ready for comparative analysis. These prove the quality of isolations used. In the next step proteins were subjected to quantitative proteomics using isobaric tandem mass tag (TMT) labeling (Fig. 3B). The proteomics of LD fraction resulted in 656 proteins and mitochondrial fraction resulted in 2,100 proteins. Both the proteomes shared 575 proteins while remaining proteins identified in the LD fraction were verified LD proteins, based on comparison with previous LD proteomes (Cermelli et al., 2006; Wang et al., 2015; Zhang et al., 2011, 2012). Using Mitocarta 2.0 (Calvo et al., 2016) and the DAVID (Huang et al., 2009b, 2009a), it was determined that LD proteome had 350 mitochondrial proteins (71.2% peptides) while mitochondrial proteome had 950 mitochondrial proteins (67.9% peptides). It was observed that there were 331 mitochondrial proteins in common between both proteomes (Fig. 3C, left). Interestingly, there were 19 mitochondrial proteins in the LD proteome which were not present in the mitochondrial proteome (Table 1). Initial proteomic analysis showed that LDAM was co-isolated with LDs and LD proteome largely contained mitochondrial proteins. So, LD proteome contained LDAM proteome and mitochondrial proteome as CM proteome. To increase the reliability, analysis threshold was set at 5 peptides. This reduced the CM proteome to 770 proteins and the LD proteome to 238 proteins (Fig. 3C, right). Of the 238 proteins in the LD fraction and 770 proteins in the mitochondrial fraction, 165 and 513, respectively, were confirmed to be mitochondrial proteins per the Mitocarta 2.0 and DAVID.

**Table 1.**
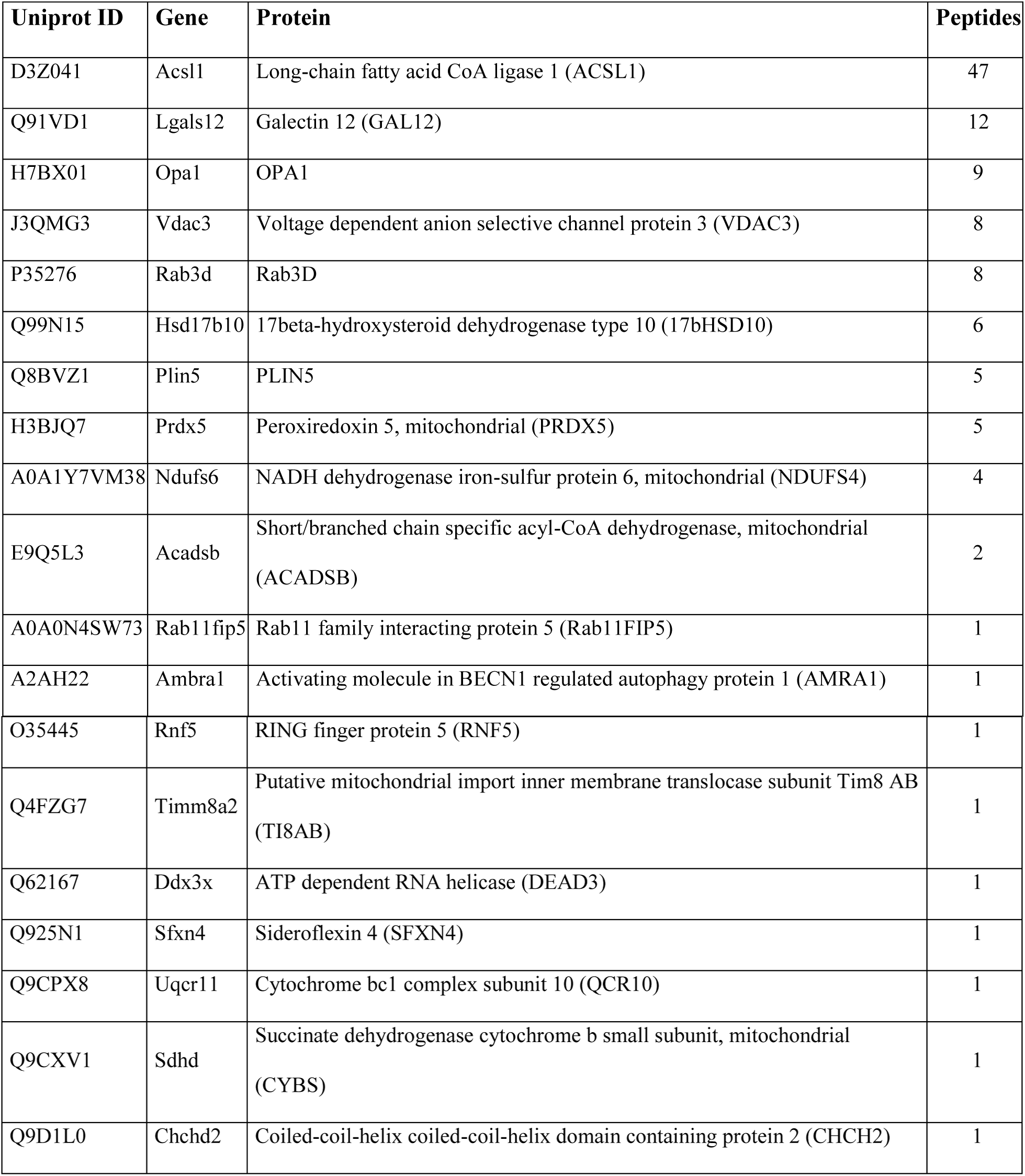
Mitochondrial Proteins in LD Proteome.

**Figure 3.**
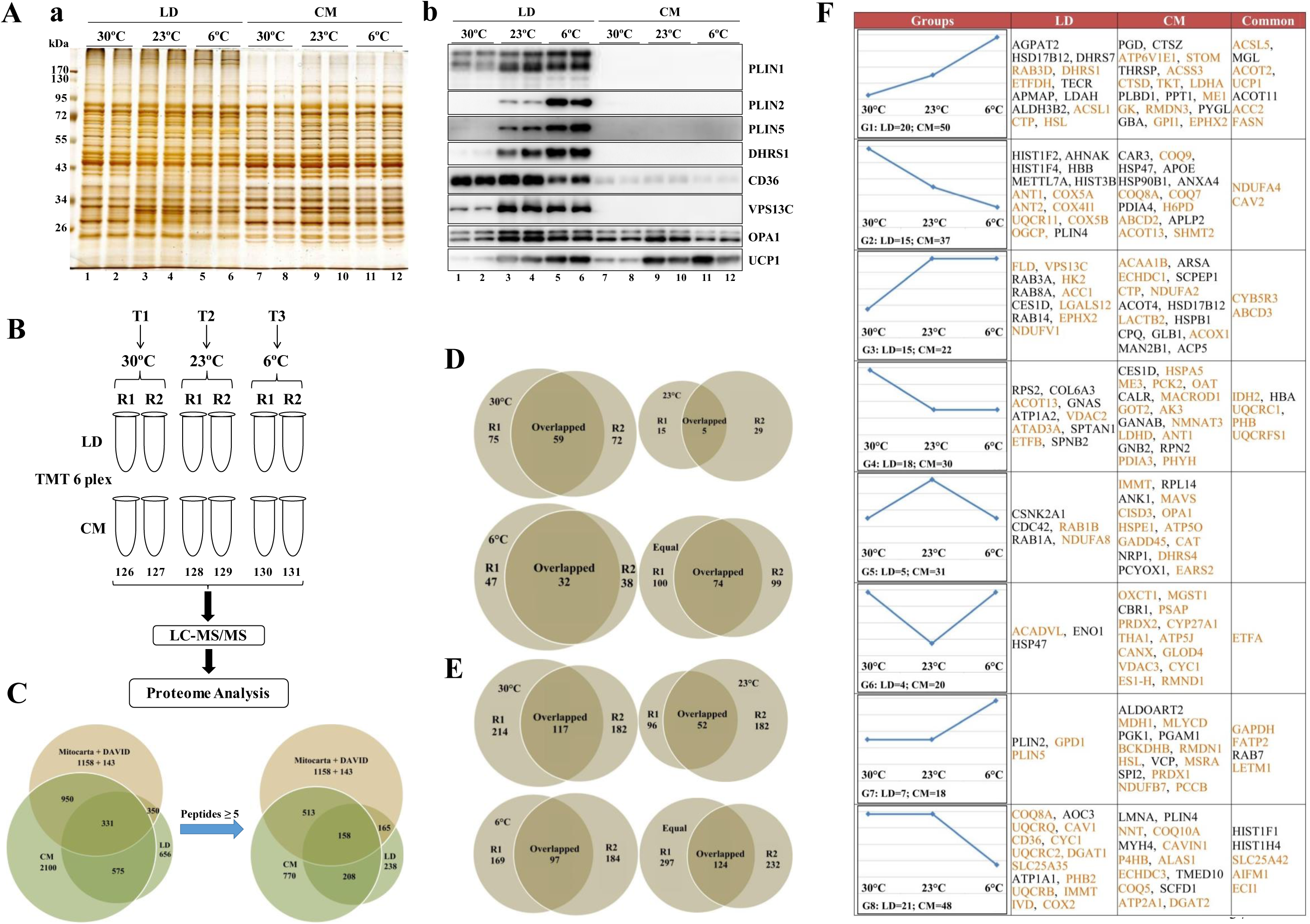
Comparative Proteomics of Lipid Droplets and Cytosolic Mitochondria of Mouse BAT from Three Temperatures. Eight-week-old male C57BL/6 mice (eight mice per group) raised at 23°C were transferred to incubators at 30°C, 23°C, or 6°C for one month. Then, their interscapular brown adipose tissue (BAT) was collected. LD and mitochondrial fractions were isolated, their proteins extracted, and analyzed by quantitative proteomics with isobaric labeling. Different bioinformatics tools and techniques were employed for the analysis. **A** Isolated BAT LDs and CM from different temperatures (30°C, 23°C, and 6°C) were analyzed by silver staining (left panel), and Western blotting (right panel). **B** Flowchart illustrating the experiment design for comparative proteomics of BAT LDs and mitochondria. **C** Venn diagram comparing LD and CM proteomes including Mitocarta 2.0 and DAVID web-service for mapping of mitochondrial proteins. A threshold of 5 peptides was set for reliable analysis. **D/E** Comparison of overlapped proteins in the LD (D) and CM (E) proteomes from R1 and R2 mice groups demonstrating proteins with high expression at 30°C, 23°C, and 6°C. The ‘Equal’ refers to the proteins which did not change in response to temperature. Only overlapping proteins were considered in the further analysis. **F** The proteins with reproducible patterns were sorted mathematically into different groups. Only proteins with significant changes in expression rates are shown (Detailed analysis can be found in Table S1). Mitochondrial proteins compared against Mitocarta 2.0 and DAVID web-service are colored in brown. Many proteins found in LD proteome belong to mitochondria. Only a few proteins showed similar expression trends in both LD and CM proteomes while most proteins showed different behaviors.

Temperature-based clustering was performed between R1 and R2 groups at each temperature to identify proteins with reproducible results. In the LD proteome there were 59, 5, and 32 proteins with high, reproducible expression at 30°C, 23°C, and 6°C, respectively (Fig. 3D, Fig. S1, left). In the CM there were 117, 52, and 97 proteins at 30°C, 23°C, and 6°C, respectively, with high, reproducible expression (Fig. 3E, Fig. S1, right). The proteins with reproducible results were further categorized into 8 groups based on expression patterns in response to temperature (Fig. 3F, Table S2). Figure 3F shows that proteins involved in β-oxidation of fatty acids (FA), e.g. ACSL5 and MGL, were commonly increased as temperature decreased. The proteins FASN and ACC2 involved in FA synthesis, also showed similar behavior (Fig. 3F, G1). UCP1 expression was also elevated at decreased temperature with the highest expression at 6°C in both LD and CM proteomes (Fig. 3F, G1). In contrast, NDUFA4, now considered to be part of the ETC C-IV (Rahman & Rahman, 2018), and CAV2 showed reduced expression with decreasing temperature with the lowest expression at 6°C in both proteomes (Fig. 3F, G2). This suggests that proteins in groups G1 and G2 are important temperature-sensitive proteins in BATs.

Proteomic analysis also showed that VPS13C (Fig. 3F, G3) and PLIN5 (Fig. 3F, G7), which were previously reported to be involved in the LD activity and in connection between organelles including mitochondria (Kumar et al., 2018; Wang et al., 2011), had high expression rates only in the LD at 6°C, suggesting that these proteins are also temperature-sensitive. The expression of other proteins, including CES1D, HSP47, and CYC1, had opposite responses between LD and CM with temperature change. It was also observed that most Rabs and histone proteins showed temperature-induced changes in expression only in LD. Together these results suggest that LD proteome has unique features and bear different protein expression patterns from CM.

### Lipid Droplet-Anchored Mitochondria Are More Sensitive to Cold

To further elaborate changes from protein to pathway level and get most out of proteomics data, an equation was developed to calculate relative strength of expression (RSE) values as:

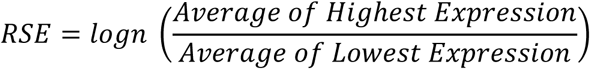

RSE values helped to rectify strength of expression of each protein in each proteome relative to different conditions, in this case temperature, in unity. It provided a simple approach to compare changes in protein expressions in LDAM and CM proteomes. Initially distribution of mitochondrial proteins was observed by clustering through Wikipathway, KEGG, Reactome databases, and Cytoscape tool (Fig. 4A). The proteins were sorted on the basis of their RSE values such that the proteins with RSE value less than 0.2 were considered with no change in expression. The number of proteins in each pathway at each RSE level were counted to determine which pathway was more affected with temperature change. It was observed that in terms of numbers most proteins with high RSE belonged to β-oxidation pathway (Fig. 4A). Later, protein clusters in different pathways were projected to heatmaps for an overview of changes (Fig. 4B). The pathway-based heatmaps showed that in comparison to CM (Fig. 4B right, Table S4), LDAM (Fig. 4B left, Table S3) showed exclusively higher expression for proteins related to β-oxidation. As previous studies of BAT have provided useful information regarding the activation of BATs at different temperatures, LDAM and CM proteomes were also compared at pathway level with previously published BAT proteomes (Table 2). It was observed that some expression patterns were common across the studies however, the LD preparation in this study demonstrated some marked differences with other LD preparations (Table 2).

**Table 2.** Pathways of LD (LDAM) and CM Proteomes.

| Proteome | LD (LDAM) |  |  | CM |  |  | PDM | CM | BAT | BAT(HFD) | BAT | hBAT |
| --- | --- | --- | --- | --- | --- | --- | --- | --- | --- | --- | --- | --- |
| Pathway | 30°C | 23°C | 6°C | 30°C | 23°C | 6°C | 22°C (1) |  | 4°C (2) | 22°C (3) | 4°C (4) | (5) |
| FA β-Oxidation | ~ | ~ | ↑ | ~ | ↑ | ↑ | ↓ | ↑ | ↑ | ↑ | ↑ | ↑ |
| TAG Synthesis | ~ | ~ | ↑ | ~ | ~ | ↑ | ↑ |  | ↑ | ↓ | ↓ |  |
| Glycolysis | ~ | ~ | ~ | ~ | ~ | ↑ |  |  | ↑ | ↑ |  |  |
| TCA Cycle | ↑ | ~ | ~ | ↑ | ↑ | ↑ | ↑ | ↓ | ↑ | ↓ | ↑ |  |
| Uncoupling Protein | ~ | ~ | ↑ | ~ | ~ | ↑ |  |  | ↑ | ↑ | ↑ | ↑ |
| ATP Synthesis | ~ | ~ | ~ | ~ | ↑ | ↑ | ↑ | ↓ |  |  | ↓ | ↓ |
| Complex I | ↑ | ~ | ~ | ↑ | ↑ | ↑ | ~ | ~ | ↑ |  | ↑ |  |
| Complex III | ↑ | ~ | ~ | ↑ | ~ | ~ | ~ | ~ | ↑ |  | ↑ |  |
| Complex IV | ↑ | ~ | ~ | ↑ | ↑ | ~ | ↑ | ↓ | ↑ |  | ↑ |  |
| Glycogenolysis | ~ | ~ | ~ | ↑ | ~ | ↑ |  |  |  |  |  |  |
| Lactate Pathway | ~ | ~ | ↑ | ↑ | ~ | ↑ |  |  |  |  |  |  |
| Glyceroneogenesis | ~ | ~ | ↑ | ↑ | ~ | ~ |  |  |  |  | ↓ |  |
| Pentose Phosphate Pathway | ~ | ~ | ~ | ~ | ↑ | ↑ |  |  |  |  |  |  |
| Malic Enzyme Based NADPH Production | ~ | ~ | ~ | ↑ | ~ | ↑ |  |  | ↑ |  |  |  |

↑ means up-regulated and ↓ means down-regulated ~ means that the expression did not show significant change as described by respective studies. LD, lipid droplet; LDAM, lipid droplet-anchored mitochondria; C, cytoplasmic mitochondria; PDM, peridroplet mitochondria; HFD, high fat diet; hBAT, human BAT. (1) = (Benador et al., 2018), (2) = (Sanchez-Gurmaches et al., 2017), (3) = (Li et al., 2015), (4) = (Forner et al., 2009), (5) = (Müller et al., 2016)

**Figure 4.**
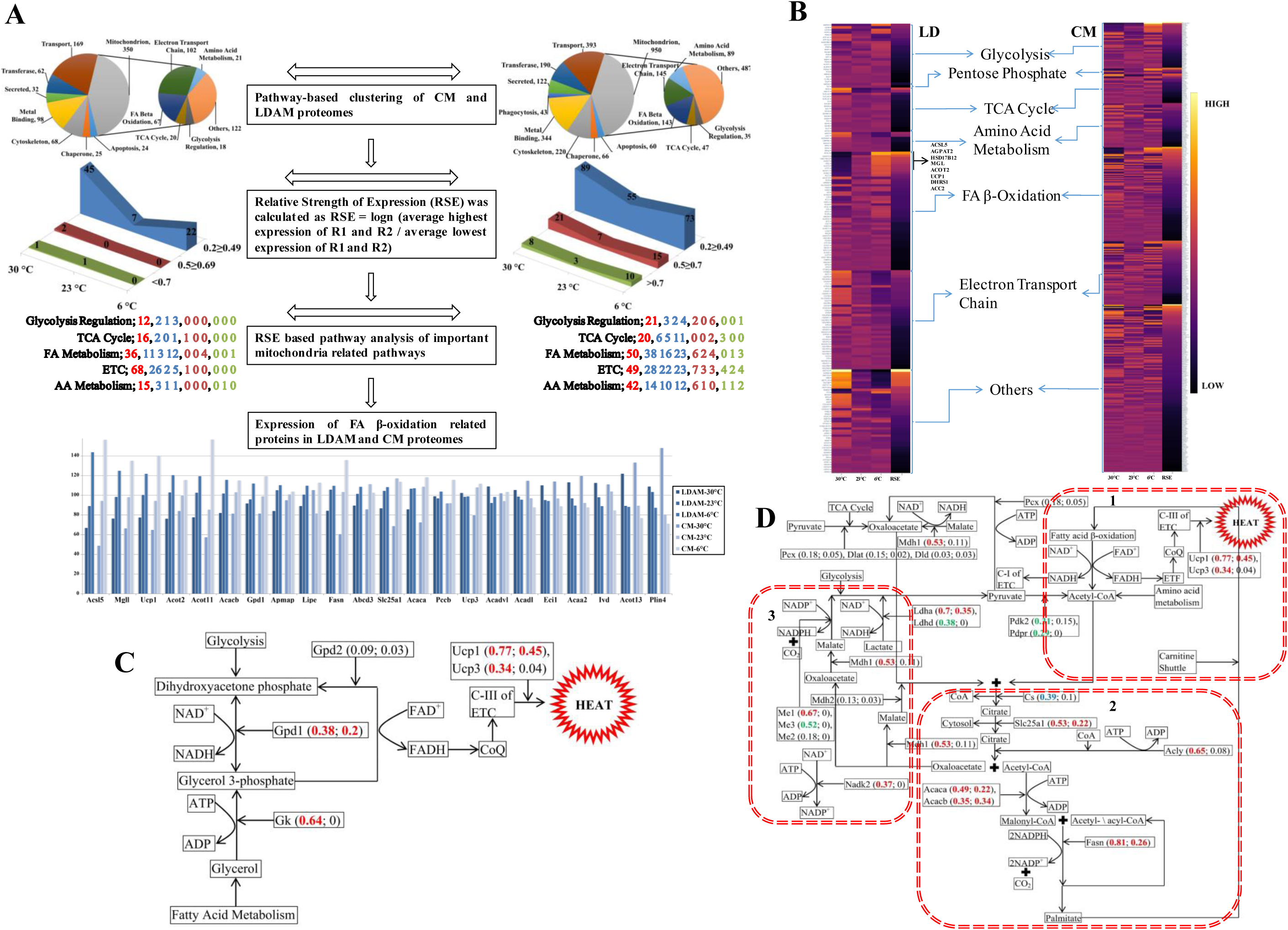
Increased Fatty Acid β-Oxidation in Lipid Droplet-anchored Mitochondria at Low Temperature. **A** Reanalysis of the proteome was performed to observe pathway level differences between LDAM and CM. Keywords based clustering was performed for whole proteome including mitochondrial pathways primarily focusing on glycolysis regulation, TCA Cycle, amino acid metabolism, ETC, and fatty acid β-oxidation. RSE values were calculated for mitochondrial pathway proteins. The proteins with RSE values less than 0.2 were considered as proteins showing no change in expression with change in temperature while higher RSE values representing higher expressions. RSE values helped in 1-dimensional (1D) analysis of expression change with respect to temperature. After RSE analysis, number of proteins at each temperature in each pathway were re-determined. The colors of numbers are according to colors of peaks for each RSE range except red color which represents proteins with no change in expression. β-oxidation proteins with significant expression change in LDAM and CM are shown. **B** Heatmaps for different mitochondrial pathways in the LD proteome and CM proteome. RSE values are mapped in the last column of each heatmap. It is easier to identify changes in expression across temperatures through RSE values; the lighter shading indicates a high expression change while darker shading indicates less expression change. The heatmap analysis demonstrated that mitochondrial pathways in LDAM responded differently from CM, with the exception of β-oxidation. **C** Mechanism for glycerol 3-phosphate (G3P) shuttle is shown. G3P shuttle transports NADH from cytoplasm to mitochondria in the form of FADH. **D** Pseudo-futile cycle is shown. When put together, RSE based deep pathway analysis shows that some sort of pseudo-futile cycle (can be termed as HEAT cycle) was active at low temperature in BAT. Pyruvate, acetyl-CoA, and citrate are leading substrates for pseudo-futile cycle fueled by FAs. The cycle can be divided into three major parts as described with dotted red boxes: 1) FA breakdown for heat production, 2) utilization of excess acetyl-CoA for FA synthesis, and 3) NADPH production as energy source for FA synthesis. **C/D** RSE values are displayed along with the proteins as protein name (CM; LDAM). Green=30°C, Blue=23°C, Red=6°C

Next, RSE-based pathway diagrams were drawn. It was observed that almost no proteins in the glycogenolysis, glycolysis, gluco/glycero/neogenesis, or pentose phosphate pathways showed significant expression changes in the LD proteome (Fig. S2). Only HK2 protein, which converts glucose to glucose 6-phosphate, showed high RSE at 23°C and GAPDH, which performs reversible catalysis of glyceraldehyde 3-phosphate to 1, 3-bisphosphoglycerate, showed a high RSE at 6°C in the LD proteome. In contrast, it was observed that with few exceptions, the CM proteome consistently showed high RSE values at 6°C for almost all proteins of glycogenolysis, glycolysis (Fig. S2), and pentose phosphate pathways (Fig. S3). Proteins of gluco/glycero/neogenesis had an elevated RSE at 30°C and the PYGL protein involved in glycogenolysis, also had a high RSE at 30°C in the CM proteome. Similarly, there were almost no proteins in the TCA cycle which had significant changes in the LD proteome except for IDH2, which had a high RSE at 30°C (Fig. S3). In the CM proteome there were differential changes in the expression of proteins in TCA cycle with some showing high RSE at the high temperature and others showing high RSE at the lower temperature (Fig. S3). There was a similarly diverse response to temperature change among ETC proteins in both the LD and CM proteomes. At 6°C no proteins involved in ATP production had high RSE in LD proteome while some did in the CM proteome. Further, in LD proteome, C-I, C-III, and C-IV proteins had elevated RSE values at 30°C and no protein had high RSE at 6°C (Fig. 4B, Table S3 and S4). The UCP3 protein had a high RSE in the CM at 6°C but had no significant changes in the LD proteome. Several proteins involved in FA β-oxidation had similar expression patterns between LD and CM. For example, HSL, MGL, ACSL5, ACOT2, and ACOT11 proteins had high RSE values at 6°C in both LD and CM (Fig. 4B, Table S3 and S4). While paradoxically, as described by Sanches-Gurmaches et. al. (Sanchez-Gurmaches et al., 2017), it was observed that proteins involved in FA synthesis, ACC2 and FASN also showed high RSE at low temperature in both proteomes. Other FA β-oxidation proteins showed different expression patterns. Overall, through proteomics study, it was observed that at lower temperature LD proteome showed higher RSE values for FA β-oxidation leading to uncoupling. Since LD fraction contained mitochondria (LDAM), the data from LD proteome suggest that the LDAM are more sensitive to cold than CM.

Further with deep pathway analysis and projection of RSE values from LD and CM proteomes onto pathway diagrams suggested that there might be two shuttle systems active at low temperature i.e. glycerol 3-phosphate shuttle (in both LD and CM) (Fig. 4C) and citrate/malate shuttle (only in CM) (Fig. 4D). It also suggested that there might be some kind of pseudo-futile cycle active in CM where degraded FAs in the form of acetyl-CoA were recycled possibly fueled by ME1 and NADK2 provided NADPH with the expense of ATP (Fig. 4D). This pseudo-futile cycle (can be termed as HEAT cycle) could be divided into three major parts as shown in red dotted lines in the Figure 4D. The first part is the UCP1 based burning of free FA by breakdown into acetyl-CoA. The excess amounts of acetyl-CoA react with oxaloacetate to form acetate which can then be shunted towards FA recycling machinery through citrate/malate shuttle. In the second part, for each acetyl-CoA, to make it part of FA chain, 2 NADPH are utilized. The third part shows malic enzyme based pathway which may provide energy for recycling of FAs. For this purpose, NADK2 protein may convert NAD^+^ into NADP^+^ with the expense of ATP while these NADP^+^ might be then converted to NADPH through malic enzyme utilizing energy released from NADH. The recycled FAs can then again be shunted towards UCP1 based burning for heat production. Currently, this cycle is called pseudo-futile cycle because it is not known that which acetyl-CoA are always used for FA recycling.

## Discussion

The interaction between mitochondria and LDs plays a vital role in thermogenesis, ectopic lipid storage, and the browning of white adipose tissue. Previous studies have shed some light on these processes but have focused on regulation events (Barneda et al., 2013; Wang et al., 2011), which do not illustrate how the two organelles act to utilize FAs efficiently and produce heat instantly. In our previous study of LDs isolated from rhesus monkey, we found that LDs anchored a subpopulation of mitochondria (LDAM) in all oxidative tissues examined, including BAT, skeletal muscle, and heart muscle, suggesting the phenomenon is essential for efficient energy utilization (Cui et al., 2019)

Previous studies found that cold-induced activation of brown adipocytes occurs through β-adrenergic signaling, leading to transcriptional regulation, including the upregulation of mitochondrial biogenesis factor PGC1α (Harms and Seale, 2013). However, exposure of a mammal to cold requires an immediate response to maintain homeostasis. Another study identified that the lipolysis of LD TAGs through β-adrenergic signal-activated TAG lipase is the first response, followed by FFA-induced mitochondrial fission and activation (Wikstrom et al., 2014). To generate heat instantly, brown adipocytes also have to transport FFAs liberated by TAG lipolysis into mitochondria for β-oxidation and induce uncoupling as well. Therefore, this multi-step process must be tightly connected, similar to the ETC, in order to produce heat immediately. The ideal scenario is that the source organelle (i.e. the LD) is in direct contact with the effector organelle (i.e. the mitochondrion), permitting rapid transfer of FFAs as fuel. A permanent physical and functional coupling between the two organelles would allow for an instant response to thermal challenge. The long-term response would involve transcriptional regulation increasing the number and activity of mitochondria.

Physical contact between LDs and mitochondria in these tissues may be in two formats, permanent and dynamic. In the current understanding, the contact between cellular organelles is dynamic, fitting a “kiss-and-run” model. Based on the discovery of Rab proteins on LDs and other early studies, researchers proposed the hypothesis of multirecognition sites in which LDs interact with almost all other membrane-bound organelles (Zehmer et al., 2009). Many subsequent studies have provided experimental support for this hypothesis (Benador et al., 2018; Pu et al., 2011; Szymanski et al., 2007; Valm et al., 2017). In at least three oxidative tissues LDs and mitochondria form a tight contact that appears to be a permanent complex. Therefore, it is proposed that in oxidative tissues the two organelles (LD and mitochondrion) may be permanently bound in a complex to increase metabolic efficiency, which supports the argument that some proteins form a rivet structure to hold two organelles (Iacovache et al., 2010). As if the rivet structure could be a channel, it may even transport FFAs from LDs to mitochondria, avoiding inefficient diffusion of hydrophobic FFAs through the aqueous cytoplasm.

## Acknowledgments

The authors thank Dr. John Zehmer for his critical reading and useful suggestions. We would like to thank Mrs. Yan Teng (Center for Biological Imaging, IBP, CAS) for her help of taking and analyzing confocal images, Wei Zhao for making Cryo-EM samples and taking the pictures, Mrs. Junfeng Hao for her help of making H&E and IHC samples, Jifeng Wang for his technical supports of comparative proteomics, Xudong Zhao and Xiaofei Guo for offering the facility equipment support. This work was supported by the National Key R&D Program of China (Grant No. 2018YFA0800700, 2016YFA0500100, and 2018YFA0800900), National Natural Science Foundation of China (Grant No. 91954108, 91857201, 31671402, 31671233, 31701018 and U1702288). This work was also supported by the “Personalized Medicines—— Molecular Signature-based Drug Discovery and Development”, Strategic Priority Research Program of the Chinese Academy of Sciences, Grant No. XDA12040218.

## Abbreviations

LD: Lipid Droplet
LDAM: LD-Anchored Mitochondria
CM: Cytoplasmic Mitochondria
FA: Fatty Acids
FFA: Free FA
BAT: Brown Adipose Tissues
RSE: Relative Strength of Expression
PDM: Peri-Droplet Mitochondria
TAG: Triacylglycerol
PLIN: Perilipin
WB: Western Blotting
TEM: Transmission Electron Microscopy
ETC: Electron Transport Chain
TMT: Tandem Mass Tag
TCA: Tricarboxylic Acid
H&E: staining Haemotoxylin and Eosin staining
DIC: Differential interference contrast
EM: Electron Microscope;

## Data availability

The mass spectrometry proteomics data have been deposited to the ProteomeXchange Consortium (http://proteomecentral.proteomexchange.org) via the PRIDE partner repository (Perez-Riverol et al., 2019) with the dataset identifier PXD018393.

## Author Contributions

P.L. conceived the project. L.C., M.A.M., and S.Z carried out experiments and data analysis. Manuscript was written by L.C., M.A.M., S.Z, and P.L.

## Competing Interests

The authors declare no competing financial interests.

